# Mapping Autosomal Recessive Intellectual Disability: Combined Microarray and Exome Sequencing Identifies 26 Novel Candidate Genes in 192 Consanguineous Families

**DOI:** 10.1101/092346

**Authors:** Ricardo Harripaul, Nasim Vasli, Anna Mikhailov, Muhammad Arshad Rafiq, Kirti Mittal, Christian Windpassinger, Taimoor I. Sheikh, Abdul Noor, Huda Mahmood, Samantha Downey, Maneesha Johnson, Kayla Vleuten, Lauren Bell, Muhammad Ilyas, Falak Sher Khan, Valeed Khan, Mohammad Moradi, Muhammad Ayaz, Farooq Naeem, Abolfazl Heidari, Iltaf Ahmed, Shirin Ghadami, Zehra Agha, Sirous Zeinali, Raheel Qamar, Hossein Mozhdehipanah, Peter John, Asif Mir, Muhammad Ansar, Leon French, Muhammad Ayub, John B. Vincent

**Affiliations:** Molecular Neuropsychiatry & Development (MiND) Lab, Campbell Family Mental Health Research Institute, Centre for Addiction and Mental Health, Toronto, ON, Canada; Institute of Medical Science, University of Toronto, Toronto, ON, Canada; Dept. of Biosciences, COMSATS Institute of Information Technology, Islamabad, Pakistan; Institute of Human Genetics, Medical University of Graz, Graz, Austria; Department of Pathology and Laboratory Medicine, Mount Sinai Hospital, Toronto, ON, Canada; Department of Laboratory Medicine and Pathobiology, University of Toronto, Toronto, ON, Canada; Fleming College, Peterborough, ON, Canada; Human Molecular Genetics Lab, Department of Bioinformatics and Biotechnology, FBAS, International Islamic University, Islamabad, Pakistan; department of Biochemistry, Quaid-i-Azam University, Islamabad, Pakistan; Qazvin University of Medical Science, Qazvin, Iran; Lahore Institute of Research & Development, Lahore, Pakistan; Department of Psychiatry, Queen’s University, Kingston, ON, Canada; Division of Hematology/Oncology, Hospital for Sick Children, Toronto, ON, Canada; Atta-ur-Rehman School of Applied Biosciences (ASAB), National University of Sciences and Technology (NUST), Islamabad, Pakistan; Department of Molecular Medicine, Biotechnology Research Center, Pasteur Institute of Iran, Tehran, Iran; Department of Biochemistry, Al-Nafees Medical College, Isra University, Islamabad, Pakistan; Department of Neurology, Qazvin University of Medical Sciences, Qazvin, Iran; Computational Neurobiology Lab, Campbell Family Mental Health Research Institute, Centre for Addiction and Mental Health, Toronto, ON, Canada; Department of Psychiatry, University of Toronto, Toronto, ON, Canada

## Abstract

Approximately 1% of the global population is affected by intellectual disability (ID), and the majority receive no molecular diagnosis. Previous studies have indicated high levels of genetic heterogeneity, with estimates of more than 2500 autosomal ID genes, the majority of which are autosomal recessive (AR). Here, we combined microarray genotyping, homozygosity-by-descent (HBD) mapping, copy number variation (CNV) analysis, and whole exome sequencing (WES) to identify disease genes/mutations in 192 multiplex Pakistani and Iranian consanguineous families with non-syndromic ID. We identified definite or candidate mutations (or CNVs) in 51% of families in 72 different genes, including 26 not previously reported for ARID. The new ARID genes include nine with loss-of-function mutations *(ABI2, MAPK8, MPDZ, PIDD1, SLAIN1, TBC1D23, TRAPPC6B, UBA7,* and *USP44),* and missense mutations include the first reports of variants in *BDNF* or *TET1* associated with ID. The genes identified also showed overlap with *de novo* gene sets for other neuropsychiatric disorders. Transcriptional studies showed prominent expression in the prenatal brain. The high yield of AR mutations for ID indicated that this approach has excellent clinical potential and should inform clinical diagnostics, including clinical whole exome and genome sequencing, for populations in which consanguinity is common. As with other AR disorders, the relevance will also apply to outbred populations.

## Introduction

Approximately 1% of the global population is affected by intellectual disability (ID)^1^, which can have a devastating effect on the lives of the affected individuals and their families and is a major challenge at the clinical level. Genetic factors are involved in the aetiology of 25-50% of ID cases^2^. The clinical presentation and aetiology of ID are complex and highly heterogeneous, thus leading to a poor rate of molecular diagnosis and inadequate clinical management and counselling.

ID can be divided into two groups: syndromic (S) ID, in which comorbid illness or physical features are present, and nonsyndromic (NS) ID, in which no such comorbidities are present. Of ~700 known ID genes (S and NS), fewer than 50 genes are mutated in NS autosomal recessive ID (NS-ARID)^2^. X-linked ID may account for only 10-12 % of ID cases^3^. Dominant autosomal variants occurring *de novo* may contribute to a large proportion of sporadic cases, particularly in outbred populations^4–6^. Autosomal recessive (AR) variants also play a significant role in ID, because recessive variants can remain in the population in heterozygous form. Estimates have suggested that there may be more than 2500 autosomal ID genes in total—the majority being recessive^7^. In populations with high levels of consanguinity, most ID-causing mutations are recessive^7^. Even in outbred populations, 13-24% of ID cases have been estimated to be due to autosomal recessive causes^7^.

Despite an increased diagnostic yield among those who receive testing^2^, currently, the majority of individuals with ID receive no molecular diagnosis^8^––a shortcoming that can affect health and lifespan. Recent studies have indicated that the median age of death is 13 years younger for males with ID and 20 years younger for females with ID than in the general population^9^. Although advances in genotyping and sequencing technology have accelerated the rate of gene discovery for ID^10^, the majority of ID genes remain undetected. However, large-scale ID family studies are making significant inroads^11^. Homozygosity mapping has been proven to be an effective method for gene identification in consanguineous populations^11–17^. Consanguineous marriages lead to a marked increase in the frequency of severe recessive disorders^18^. Collectively, countries with levels of consanguinity higher than 10%, mainly in Africa, the Middle East and South Asia^19, 20^, represent a population of ~1.3 billion. In Pakistan, ~62.7% of the population engages in consanguineous marriages, of which ~80.4% are first-cousin marriages^21^. The rate of consanguineous marriages in Iran is estimated at 40%^22^. Here, we present a study of multiplex ID families from Pakistan (N=176) and Iran (N=16) using microarray-based genotyping to identify autozygous regions (HBD shared by affected family members), coupled with whole exome sequencing (WES) to identify causal variants (see Figure 1 for workflow). In total, we identified single candidate genes/variants in 88 families (50% were loss-of-function (LoF) mutations), and ten candidate pathogenic genomic variants (CNVs) in nine families.

**Figure 1.**
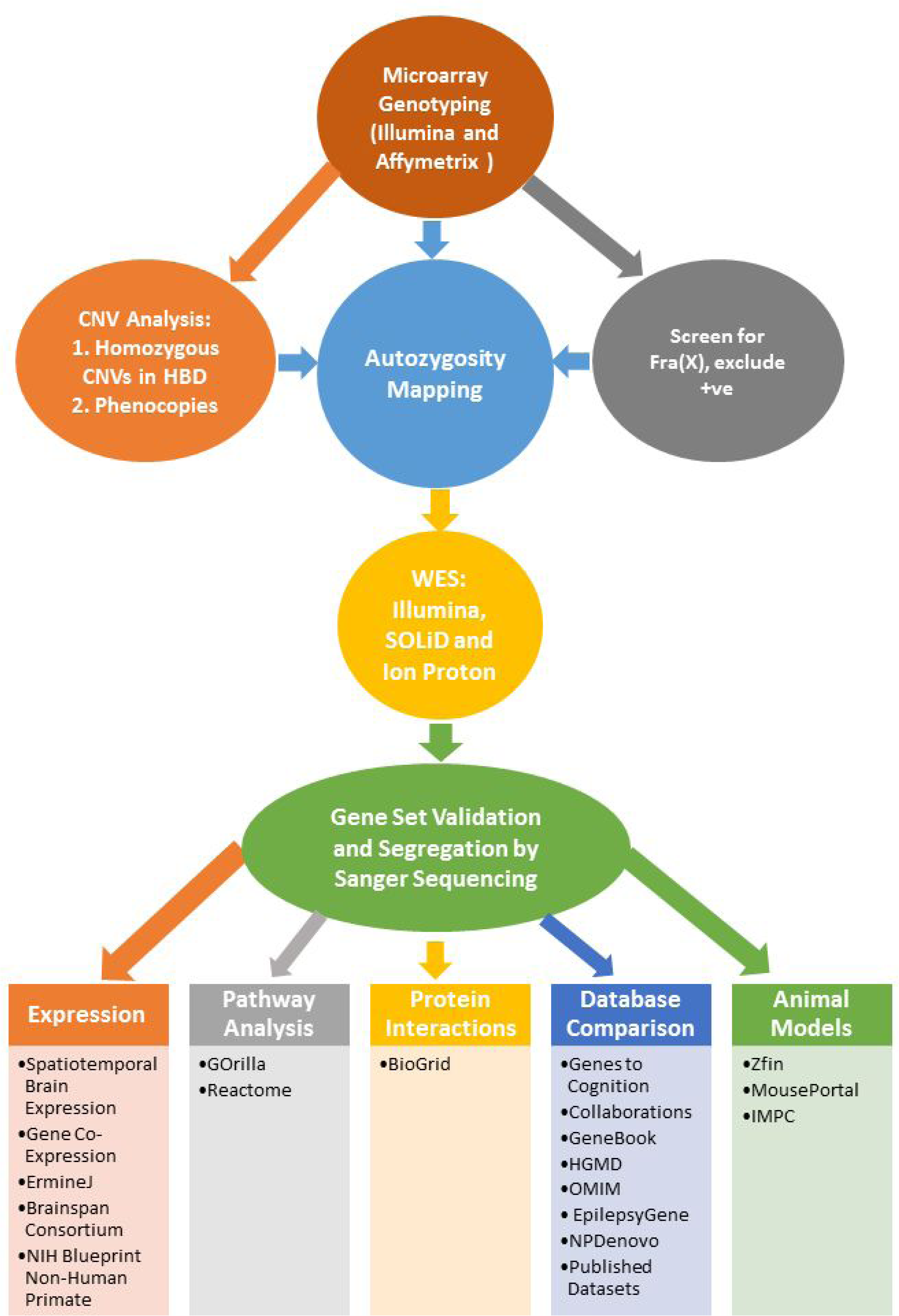
Microarray and exome sequencing work flow.

## Methods

### Family recruitment

Institutional research ethics board consent was given for the study by the Centre for Addiction & Mental Health, Toronto, as well as by the institutes at the recruiting sites (details in the Supplementary Methods). Families were recruited on the basis of diagnosis of ID in more than one individual (or ID and/or learning disability for N=13 families; or both ID and psychosis within the family, N=5) but with no obvious dysmorphic features or comorbidities and with parental consanguinity. Typically marriages were first-or second-cousin marriages; however, in a number of families, the exact relationship between the parents could not be established, but the marriage was within the same clan or caste. Written informed consent was obtained from all participants. Blood was drawn, and genomic DNA was extracted by standard methods. Cases of fragile X (tested using established methods^23, 24^), Down’s syndrome and other clearly recognizable syndromes were excluded. Summary statistics for the families are given in Table 1.

**Table 1.**
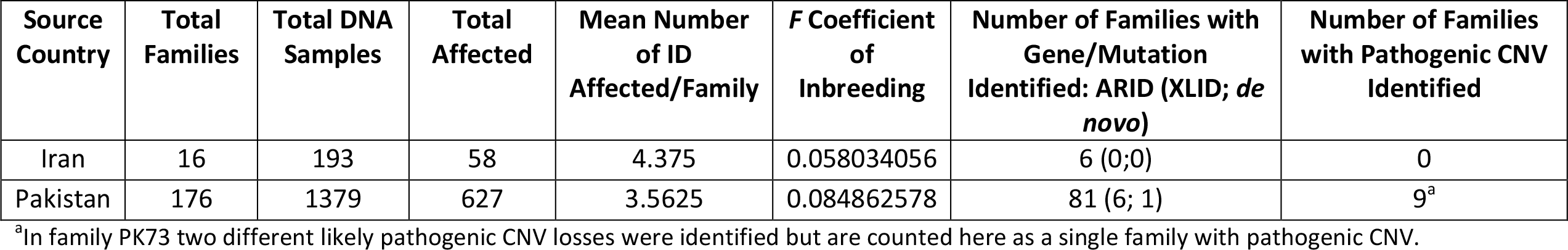
Summary statistics for the Iranian and Pakistani family cohorts.

### Autozygosity Mapping

Autozygosity mapping was performed using microarray data from the Illumina Human CoreExome, Affymetrix Mapping 500K *Nsp*I, Affymetrix CytoScan HD and Affymetrix SNP 5.0 or 6.0. All genotyping was performed according to the manufacturer’s protocol, and data were processed using either the Affymetrix Genotyping Console (500K and 5.0), Affymetrix ChAS Software Suite (CytoScan HD and SNP 6.0) or the Illumina GenomeStudio platform (Illumina CoreExome). All data were exported to PLINK format for analysis using HomozygosityMapper^25^ and FSuite^26^. Details are provided in the Supplementary Methods.

### CNV Analysis

CNV analysis was performed to identify homozygous CNVs in HBD regions or heterozygous CNVs that could indicate cases of intra-familial genetic heterogeneity, which were then excluded from the HBD/WES analyses (details in the Supplementary Methods).

### Whole Exome Sequencing (WES)

WES was performed for one or more affected member from each family, using sequencing facilities at CAMH, Toronto. Three different next-generation sequencing platforms were used across the study, taking advantage of newer platforms as they became available for research at CAMH (SOLiD 5500 platform (Life Technologies) for 49 families (51 individuals), using a protocol reported previously^27^; Ion Proton platform/Ion Ampliseq™ Exome kit (Life Technologies) for 49 individuals from 30 families; Illumina HiSeq2500 platform, using ThruPLEX DNA-seq 96D kit (Rubicon, R400407) and SureSelect XT2 Target Enrichment (Agilent Technologies) system for 150 families).

### Sequencing Alignment and Variant Calling

We used an in-house pipeline to map and call variants on the different types of sequencing data for this study. The pipeline is summarized in Supplementary Figure 1 and in the Supplementary Methods.

After putative variants were identified, Sanger sequencing was used to validate the variants and determine whether the variant segregated with the disease by testing the parents and other affected and unaffected individuals within the family.

### Gene list construction

We combined the genes listed in Supplementary Table 3 with other genes for X-linked or autosomal recessive intellectual disability/mental retardation in OMIM. This full list is available as Supplementary Table 7. This gene list was used in the pathway and gene expression analyses (details in the Supplementary Methods).

### Database searches across different neuropsychiatric and neurodevelopmental disorders and functional and animal models

We compared our variants to several databases including EpilepsyGene, Gene2Phenotype, Gene2Cognition, Schizophrenia Genebook, published disease-specific gene-sets^6^, HGMD^28^, OMIM^29^, Zfin^30^, targets of FMRP through high-throughput sequencing of RNAs by cross-linking immune-precipitation (HITS-CLIP) ^31^, and Mouseportal (http://www.sanger.ac.uk/science/collaboration/mouse-resource-portal, accessed May 2016). Searches were performed to identify genes/variants overlapping with our dataset.

## Results

Our study identified single candidate genes/variants for 88 (81 autosomal recessive, 6 X-linked, and 1 *de novo* (heterozygous)) of the 192 families (Table 2 & 3, & Supplementary Table 3a). Twenty-six of the genes identified in this cohort have not previously been reported for NS-ARID. An additional eleven genes were previously first reported from this cohort *(CC2D2A^32^; TCTN2^33^; TRAPPC9^12^; MAN1B1^13^; FBXO31^27^; METTL23^14^; FMN2^15^; DCPS^16^; HMNT^17^; NSUN2*^34^; *MBOAT7^35l^).* Most of these genes have since been reported in multiple ID families, including in outbred populations (e.g., *TRAPPC9*^36^*; MAN1B1*^37^). Likely pathogenic CNVs were identified in at least nine families (not including the VPS13B and TUSC3 deletions in AN51 and AN21, respectively).

**Table 2.** Novel candidate genes and sequence variants identified in the Pakistani and Iranian ID families. A list of all variants identified including known syndromic ARID or XLID genes identified in the cohort is given in Supplementary Table 3a; Supplementary Table 3b lists families in which between 2 and 4 variants were identified.. See Supplementary Table 3 for *in silico* predictions of the effects of the variants.

|  | Family | Gene | Genome (ExAC MAF/S. Asian MAF) | cDNA (Protein) |
| --- | --- | --- | --- | --- |
| 1 | PJ7 | *LAMC1* | Chr1:183083732A>T (0/0) | NM_002293.3:c.1088A>T (p.His363Leu) |
| 2 | PK113 | *AFF3* | Chr2:100167973C>A (0/0) | NM_002285.2:c.3644G>T (p.Gly1215Val) |
| 3<sup>a</sup> | AS61 | *ABI2* | Chr2:204245039C>T (0/0) | NM_005759.4:c.394C>T (p.Arg132*) |
| 4<sup>b</sup> | PK34 | *UBA7* | Chr3:49848458C>A (8.01E-04/5.45E-03) | NM_003335.2:c.1189G>T (p.Glu397*) |
| 5 | PJ9 | *TBC1D23* | Chr3:100037953delA (0/0) | NM_018309.4:c.1683delA (p.Glu562Argfs*3) |
| 6 | AN50 | *ZBTB11* | Chr3:101371344A>C (0/0) | NM_014415.3:c.2640T>G (p.His880Gln) |
| 7 | AS70 | *MAP3K7* | Chr6:91254333C>T (1.65E-05/1.21E-04) | NM_145331.2:c.1229G>A (p.Arg410Gln) |
| 8 | PJ2 | *SUMF2* | Chr7:56146063A>G (8.70E-06/7.73E-5) | NM_015411.2:c.740A>G (p. Tyr247Cys) |
| 9 | AS20 | *EXTL3* | Chr8:28,575,513C>G (0/0) | NM_001440.3:c.1937C>G (p. Ser646Cys) |
| 10<sup>c</sup> | AS66 | *MPDZ* | Chr9:13109992_13109995delGAAA (0/0) | NM_003829.4:c.5811_5814delTTTC (p.Phe1938Leufs*3) |
| 11 | AS22 | *MAPK8* | Chr10:49628265_49628265delC (0/0) | NM_002750.3:c.518delC (p.Arg174Glyfs*8) |
| 12 | PK70 | *TET1* | Chr10:70451328G>T (0/0) | NM_030625.2:c.6168G>T (p.Lys2056Asn) |
| 13 | IDSG19 | *DMBT1* | Chr10:124356559C>G (0/na) | NM_007329.2:c.2906C>G (p.Thr969Arg) |
| 14<sup>d</sup> | AS105 & 110 | *PIDD1* | Chr11:799453G>A (0/0) | NM_145886.3:c.2587C>T (p.Gln863*) |
| 15 | PK135 | *BDNF* | Chr11:27679747A>G (1.65E-04/1.21E-03) | NM_001709.4:c.365T>C (p.Met122Thr) |
| 16 | PK94 | *USP44* | Chr12:95927147_95927160delinsA (0/0) | NM_032147.3:c.873_886delinsT (p.Leu291Phefs*8) |
| 17 | AS23 | *SPATA13* | Chr13:24797332C>T (4.71E-05/0) | NM_001166271.1:c.265C>T (p.Arg89Trp) |
| 18 | PK68 | *SLAIN1* | Chr13:78320722_78320722delA (0/0) | NM_144595.3:c.135delA (p.Thr49Hisfs*96) |
| 19 | IDH3 | *TRAPPC6B* | Chr14:39628712G>A (1.65E-05/na) | NM_177452.3:c.124C>T (p.Arg42*) |
| 20 | PK79 | *VPS35* | Chr16:46705735T>G (0/0) | NM_018206.4:c.1406A>C (p.Gln469Pro) |
| 21 | PK91 | *SYNRG* | Chr17:35896199C>T (8.66E-06/0) | NM_007247.4:c.3548G>A (p.Arg1183His) |
| 22 | IDSG29 | *FBXO47* | Chr17:37107906G>C (0/na) | NM_001008777.2:c.544C>G (p.Arg182Gly) |
| 23 | AN53 | *SDK2* | Chr17:71431658C>T (5.82E-05/0) | NM_001144952.1:c.1126G>A (p.Gly376Ser) |
| 24 | AN48 | *CAPS* | Chr19:5914970G>A (2.17E-04/1.27E-04) | NM_004058.3:c.281G>A (p.Arg94Gln) |
| 25 | PJ6 | *GPR64* | ChrX:19055718T>C (8.45E-05/0) | NM_005756.3:c.191A>G (p.Asn64Ser) |
| 26 | PK87 | *MAGEA11* | ChrX:148796213C>T (0/0) | NM_005366.4:c.169T>C (p.Ser57Pro) |

- The homozygous nonsense mutation reported for family AS61 is located within a large HBD region shared between only two of the four affected individuals. The other two are phenotypically distinct with ID and myopathy. Support for this gene comes from functional studies, including with *Abi2* null mice<sup>96</sup>. - Two homozygotes for this variant are indicated in the ExAC South Asian population (N=8,251 individuals). N.B. affected individuals in PK34 have learning disability rather than ID. The South Asian cohort used in ExAC was part of the Pakistan Risk of Myocardial Infarction Study (PROMIS) and would not have excluded subjects in this category. - The *MPDZ* mutation reported here is not within a significant region of HBD and is present as homozygous in the affected mother and the first-born affected son. The affected brother and sister are both heterozygous, but the father is wild type homozygous, thus suggesting non paternity for the first child. - Families AS105 and AS110 both carry the same mutation in *PIDD1* and share a common haplotype; however, these families are apparently not closely related. Using Prest-plus<sup>81</sup> to estimate IBD using Maximum Likelihood Estimation, it was confirmed that the two families are separated by at least four generations, which is
comparable to relatedness in the background population, and thus AS105 and AS110 are considered unrelated. na=not applicable, because Iranian exome data are not available in the ExAC database.

**Table 3.**
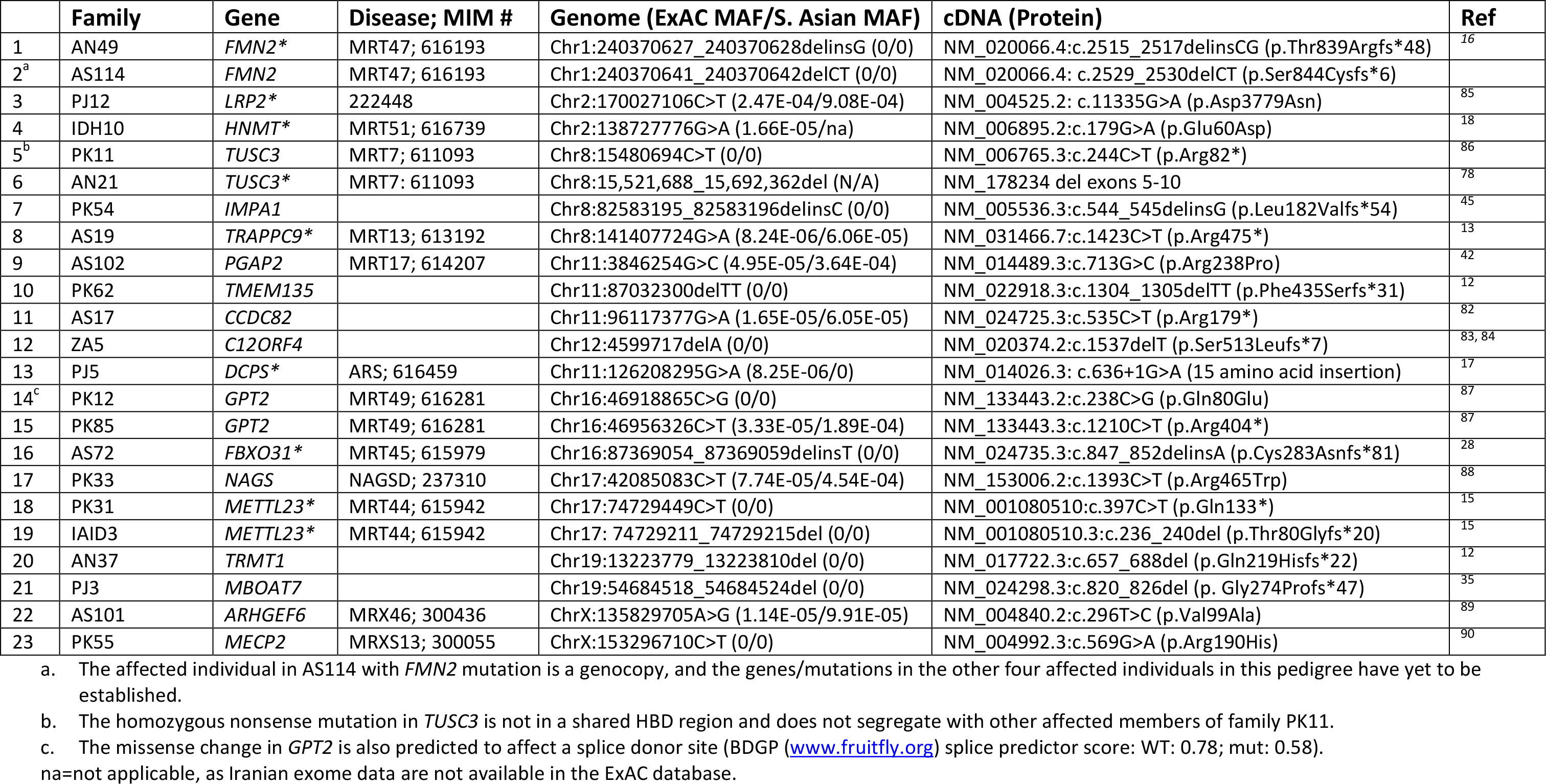
Candidate genes and sequence variants identified in the Pakistani and Iranian ID families in support of previously reported nonsyndromic ID genes. Asterisks indicate genes/variants that we have previously reported in this cohort. For subjects in whom the phenotype was clearly NS-ARID, despite prior association of the gene with S-ARID (e.g. *LRP2* and *MECP2*), we have included the variant in this list. See Table S3 for *in silico* predictions of the effects of the variants.

|  | Family | Gene | Disease; MIM # | Genome (ExAC MAF/S. Asian MAF) | cDNA (Protein) | Ref |
| --- | --- | --- | --- | --- | --- | --- |
| 1 | AN49 | *FMN2* * | MRT47; 616193 | Chr1:240370627_240370628delinsG (0/0) | NM_020066.4:c.2515_2517delinsCG (p.Thr839Argfs*48) | 16 |
| 2<sup>a</sup> | AS114 | *FMN2* | MRT47; 616193 | Chr1:240370641_240370642delCT (0/0) | NM_020066.4: c.2529_2530delCT (p.Ser844Cysfs*6) |  |
| 3 | PJ12 | *LRP2* * | 222448 | Chr2:170027106C>T (2.47E-04/9.08E-04) | NM_004525.2: c.11335G>A (p.Asp3779Asn) | 85 |
| 4 | IDH10 | *HNMT* * | MRT51; 616739 | Chr2:138727776G>A (1.66E-05/na) | NM_006895.2:c.179G>A (p.Glu60Asp) | 18 |
| 5<sup>b</sup> | PK11 | *TUSC3* | MRT7; 611093 | Chr8:15480694C>T (0/0) | NM_006765.3:c.244C>T (p.Arg82*) | 86 |
| 6 | AN21 | *TUSC3* * | MRT7; 611093 | Chr8:15,521,688_15,692,362del (N/A) | NM_178234 del exons 5-10 | 78 |
| 7 | PK54 | *IMPA1* |  | Chr8:82583195_82583196delinsC (0/0) | NM_005536.3:c.544_545delinsG (p.Leu182Valfs*54) | 45 |
| 8 | AS19 | *TRAPPC9* * | MRT13; 613192 | Chr8:141407724G>A (8.24E-06/6.06E-05) | NM_031466.7:c.1423C>T (p.Arg475*) | 13 |
| 9 | AS102 | *PGAP2* | MRT17; 614207 | Chr11:3846254G>C (4.95E-05/3.64E-04) | NM_014489.3:c.713G>C (p.Arg238Pro) | 42 |
| 10 | PK62 | *TMEM135* |  | Chr11:87032300delTT (0/0) | NM_022918.3:c.1304_1305delTT (p.Phe435Serfs*31) | 12 |
| 11 | AS17 | *CCDC82* |  | Chr11:96117377G>A (1.65E-05/6.05E-05) | NM_024725.3:c.535C>T (p.Arg179*) | 82 |
| 12 | ZA5 | *C12ORF4* |  | Chr12:4599717delA (0/0) | NM_020374.2:c.1537delT (p.Ser513Leufs*7) | 83, 84 |
| 13 | PJ5 | *DCPS* * | ARS; 616459 | Chr11:126208295G>A (8.25E-06/0) | NM_014026.3: c.636+1G>A (15 amino acid insertion) | 17 |
| 14<sup>c</sup> | PK12 | *GPT2* | MRT49; 616281 | Chr16:46918865C>G (0/0) | NM_133443.2:c.238C>G (p.Gln80Glu) | 87 |
| 15 | PK85 | *GPT2* | MRT49; 616281 | Chr16:46956326C>T (3.33E-05/1.89E-04) | NM_133443.3:c.1210C>T (p.Arg404*) | 87 |
| 16 | AS72 | *FBXO31* * | MRT45; 615979 | Chr16:87369054_87369059delinsT (0/0) | NM_024735.3:c.847_852delinsA (p.Cys283Asnfs*81) | 28 |
| 17 | PK33 | *NAGS* | NAGSD; 237310 | Chr17:42085083C>T (7.74E-05/4.54E-04) | NM_153006.2:c.1393C>T (p.Arg465Trp) | 88 |
| 18 | PK31 | *METTL23* * | MRT44; 615942 | Chr17:74729449C>T (0/0) | NM_001080510:c.397C>T (p.Gln133*) | 15 |
| 19 | IAID3 | *METTL23* * | MRT44; 615942 | Chr17: 74729211_74729215del (0/0) | NM_001080510.3:c.236_240del (p.Thr80Glyfs*20) | 15 |
| 20 | AN37 | *TRMT1* |  | Chr19:13223779_13223810del (0/0) | NM_017722.3:c.657_688del (p.Gln219Hisfs*22) | 12 |
| 21 | PJ3 | *MBOAT7* |  | Chr19:54684518_54684524del (0/0) | NM_024298.3:c.820_826del (p. Gly274Profs*47) | 35 |
| 22 | AS101 | *ARHGEF6* | MRX46; 300436 | ChrX:135829705A>G (1.14E-05/9.91E-05) | NM_004840.2:c.296T>C (p.Val99Ala) | 89 |
| 23 | PK55 | *MECP2* | MRXS13; 300055 | ChrX:153296710C>T (0/0) | NM_004992.3:c.569G>A (p.Arg190His) | 90 |

- The affected individual in AS114 with *FMN2* mutation is a genocopy, and the genes/mutations in the other four affected individuals in this pedigree have yet to be established. - The homozygous nonsense mutation in *TUSC3* is not in a shared HBD region and does not segregate with other affected members of family PK11. - The missense change in *GPT2* is also predicted to affect a splice donor site (BDGP (www.fruitfly.org) splice predictor score: WT: 0.78; mut: 0.58). - na=not applicable, as Iranian exome data are not available in the ExAC database.

### Homozygous Loss-of-Function Mutations

Homozygous truncating LoF mutations were found as single candidate variants in 43 families, including nine genes that had not been previously described in relation to NS-ARID: *ABI2, MAPK8, MPDZ, PIDD1, SLAIN1, TBC1D23, TRAPPC6B, UBA7,* and *USP44.* None of these genes have been noted as tolerant to loss of function through WES of 3,222 British adults of Pakistani origin^38^.

We identified the same nonsense mutation in *PIDD1* in two unrelated Pakistani families (ASMR105 and ASMR110; Figure 2). *PIDD1* encodes p53-Induced Death Domain Protein (PIDD; MIM 605247), and Gln863* disrupts the death domain (DD), through which PIDD1 interacts with other DD proteins such as RIP1 or CRADD/RAIDD. Truncating mutations in *CRADD* have previously been reported for NS-ARID (MRT34^39^), and thus our findings support the involvement of PIDD-related pathways in the aetiology of ID.

**Figure 2.**
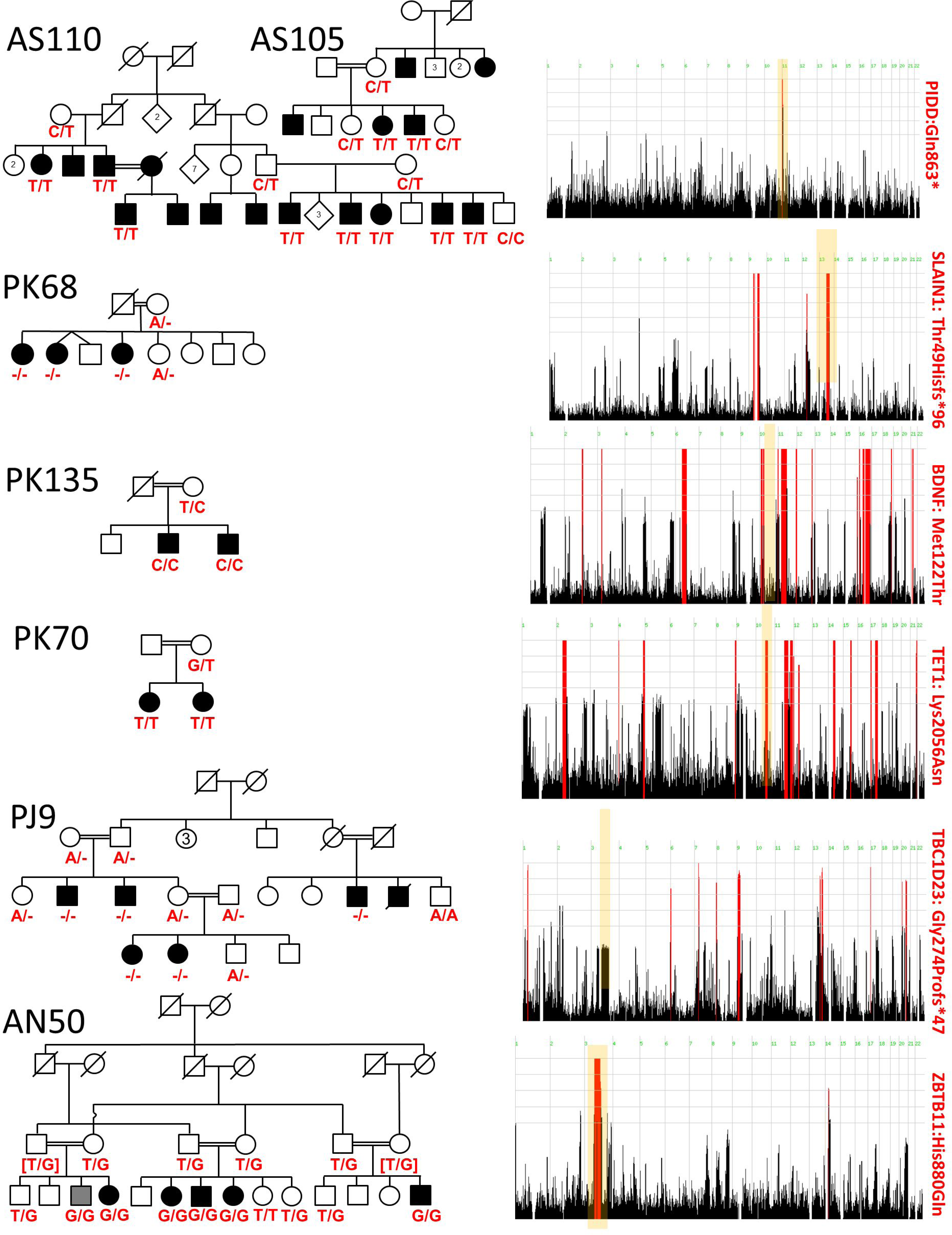
Pedigrees and HomozygosityMapper output for eight of the families. The locus harbouring the mutation is indicated in the HomozygosityMapper plots with yellow shading. HBD regions for each family, confirmed as autozygous and haploidentical, are provided in Supplementary Table 2. The gene name and mutation (at the protein level) are indicated, as well as genotypes for available family members. Additional pedigrees and HBD plots are provided in Supplementary Figure 2.

Homozygous truncating mutations in *TRAPPC9* have been reported for NS-ARID ^12,36, 40–43^; thus it is notable that we identified a nonsense mutation in *TRAPPC6B,* which encodes a second member of the same protein trafficking particle complex.

A nonsense mutation was identified in *SLAIN1* that segregated fully in family PK68 (Figure 2). *SLAIN1* and *SLAIN2* encode microtubule plus-end tracking proteins that have been shown to be crucial for normal axonal growth in developing hippocampal neurons^44^.

We report here a homozygous nonsense mutation in the gene *UBA7* in family PK34. *UBA7* encodes ubiquitin-activating enzyme 7, which is believed to be involved in the ubiquitin conjugation pathway^45^. Family PK34 is one of three families in the study that did not meet the criteria for ID and instead were reported as having learning disorders and were considered relatively high functioning. We also report that this variant is present at a relatively high frequency in the South Asian population (MAF=0.0054; ExAC database), and two homozygotes for this variant were among the control group. Thus, this variant and gene may be a risk factor for a much milder form of cognitive disability and thus may potentially be present in the control South Asian population (N>8,000) used in the ExAC database.

We also identified a LoF mutation in *TMEM135,* which has previously reported in a large Iranian NS-ARID cohort by Najmabadi et al, 2011 (in which a missense mutation, Cys228Ser, was reported)^11^, and in Wright et al, 2014 (missense)^46^, as a quantitative trait locus associated with intelligence^47^.

### Missense mutations

We identified homozygous missense variants as single candidate variants in 43 families, including 16 for which the identified genes have not previously been reported for NS-ARID *(AFF3, BDNF, CAPS, DMBT1, DUOX2, EXTL3, FBXO47, LAMC1, MAP3K7, SDK2, SPATA13, SUMF2, SYNRG, TET1, VPS35,* and *ZBTB11).* Although LoF mutations are frequently more convincing than missense mutations, a number of the homozygous missense changes reported here are of particular interest, owing to the known nature or function of the protein or the likely effect of the amino acid substitution on protein function. For instance, we report a homozygous missense change Met122Thr in the gene for brain-derived neurotrophic factor, *BDNF,* which has been implicated in many studies of neuropsychiatric disorders^48^ and is a known gene target of *MECP2,* the Rett syndrome gene.^49, 50^ This variant replaces a methionine, a large, non-polar residue with an S-methyl thioether side chain, that is fully conserved across the vertebrate lineage with threonine, a small, polar residue with a hydroxyl side chain. This variant is present with a frequency of 0.0001647 in ExAC (in South Asians MAF=0.001211) with no homozygotes and has not previously been reported in any publications. The BDNF protein is important for the survival, differentiation and development of neurons in the central nervous system (CNS). Hence, the identification of Met122Thr in connection with cognitive disability is likely to be of great interest, and functional studies of the effects of Met122Thr on BDNF function are warranted.

We detected a homozygous Lys2056Asn change in *TET1* in family PK70 that is not present in any variant or mutation databases. Methylcytosine dioxygenase TET proteins play a role in the DNA methylation process and gene activation, and TET1 is an important regulator of neuronal differentiation. TET1 has been implicated in schizophrenia (SCZ) and bipolar disorder (BD) by down-regulation of the expression of *GAD1, RELN,* and *BDNF* genes through epigenetic mechanisms in the prefrontal cortex and in the cerebellum in autism^51, 52^. Significant upregulation of *TET1* mRNA in the parietal cortex of psychosis patients has also been reported^53^. *Tet1* knockout results in impaired hippocampal neurogenesis, cognitive deficits in mice^54^ and abnormal brain morphology in zebrafish^55^. The large basic, charged, hydrophobic lysine residue at this position is conserved in mammals (or as glutamic acid in non-mammalian vertebrates) and is replaced in this variant by the small polar asparagine residue.

The homozygous His880Gln change identified in the zinc finger/BTB domain gene *ZBTB11* in family AN50 disrupts a canonical Zn^2+^-binding residue in one of the zinc fingers and is likely to result in an alteration in the specificity of DNA-binding/gene regulation. *AFF3* is an autosomal homolog of the X-linked ID gene *AFF2* (MRX-FRAXE; MIM 309547). As with *AFF2, AFF3* is also associated with a fragile site (FRA2A) in which expansion of a CGG trinucleotide repeat triggers hypermethylation and gene silencing in association with neurodevelopmental disorders^56^. The Gly1215Val variant identified here is located within a five-amino acid C-terminal motif (Gln-Gly-Leu-His-Trp) that is highly conserved across the vertebrate lineage.

In family AS70, a missense mutation was identified in exon 12 of the gene *MAP3K7.* Although missense mutations in this gene have recently been reported for frontometaphyseal dysplasia^57^ (FMD) and cardiospondylocarpofacial syndrome (CSCF syndrome)^58^, members of family AS70 have no skeletal dysplasia or obvious dysmorphic features, thus suggesting even greater pleiotropy of this gene. We note that although the Arg410Gln mutation identified is located in exon 12, which is alternatively spliced and present in both transcripts B and C but neither A nor D, all *MAP3K7* mutations reported to date for FMD or CSCF syndrome are located in canonical exons (i.e., in all four transcript variants). Furthermore, analysis of mouse RNAseq data from the ENCODE UW project (through UCSC Genome Browser) suggests that although exon 12 is expressed in many adult tissues, including the brain, it is absent from skeletal muscle.

### Families with multiple variants

For eleven additional families, between two and four putative damaging variants were identified after filtration that fulfilled the criteria and segregated in the families; however, without functional evidence in support of pathogenicity, it is not possible to narrow these variants to a single candidate (Supplementary Table 3B).

### X-linked variants

A number of the families were compatible for both AR and X-linkage, and several variants or CNVs on the X chromosome were identified, including two variants in *ATRX* and a 6.7-Mb interstitial duplication on Xp22.31-p22.2.

We report a missense mutation, Ser57Pro, in the X-chromosomal gene *MAGEA11,* segregating in family PK87. A Gln4Arg variant in MAGEA11 has recently been reported among a cohort of X-linked ID families^59^. MAGEA11 shows protein interaction with TRMT1—also reported here for NS-ARID—as determined by yeast-two-hybrid analysis^60–62^.

In another large multiplex and multi-branch family, the missense mutation Arg190His of the X-linked *MECP2* gene is present in hemizygous form in a single male with mild ID and in heterozygous form in several females with mild ID or mild ID with psychosis or depression. Interestingly, for the female heterozygotes with ID, all appeared cognitively normal until the age of ~9 years, when cognitive regression started, and for the single male hemizygote, cognitive regression began much earlier, at <5 years of age. The family also has several males with SCZ, with onset at ~18 years but without cognitive regression amounting to ID, who are wild type for this variant (see Supplementary Figure 2). The Arg190 residue is a critical DNA-binding amino acid within an AT-hook domain, and a *de novo* mutation at this residue, Arg190Cys, has been reported in a SCZ patient^63^.

### Genes with ‘hits’ in multiple families

Confidence in gene discovery relies on the identification of multiple affected families with mutations in the same gene and/or replication in further studies. However, owing to the anticipated high degree of genetic heterogeneity in NS-ARID, large sample sizes are typically required to include multiple families with mutations in the same gene. Comparison across different studies is thus vital. Of the ARID genes reported here for the first time, several are conspicuous because of the presence of mutations in multiple families. In our study, the same nonsense mutation in *PIDD1* was observed in two apparently unrelated Pakistani families. We also provide confirmatory families/mutations for recently reported ARID genes such as *C12ORF4, CCDC82, MBOAT7, IMPA1, TMEM135, PGAP2, GPT2* (2 families), *TDP2* (2 families), and *GMPPA* (Table 3, and Supplementary Table 3). In addition, mutations in the gene for ID-associated brain malformation polymicrogyria, *GPR56,* are present in three families in this study and thus may represent a relatively large proportion of ID families in these populations.

### Families with mutations in previously identified genes for metabolic or hormonal disorders

We report three mutations in the thyroid dyshormonogenesis (TDH) genes *TPO* (thyroid peroxidase), *TG* (thyroglobulin)^64^), and *DUOX2* (thyroid oxidase 2). With adequate clinical resources available in most developed countries, many of these cases would likely have been diagnosed, and in some cases, for example, those with mutations in N-acetylglutamate synthase *(NAGS), TPO, TG* and *DUOX2,* early treatment would have prevented ID development. Given the prevalence of mutations in these genes in a relatively modest number of families, it is likely that these disorders are relatively common causes of ID in populations in which consanguinity is common, but access to clinical diagnostics is poor. Mutations in *GNE* have previously been linked with sialuria (dominant; MIM 269921)—an extremely rare metabolic disorder—and Nonaka myopathy (recessive; MIM 605820), whereas here we report a homozygous missense mutation in a family with only ID and no myopathy. Although we were unable to perform biochemical analysis of this family, we anticipate that this discovery may represent a previously unreported recessive form of sialuria.

### Copy number variation (CNV) analysis

In addition to HBD analysis, the microarray data were used for CNV analysis: first, to identify possible intra-familial genetic heterogeneity with large, probably pathogenic heterozygous loss/gain CNVs, and second, to detect homozygous genic CNVs within mapped HBD regions. For the former, several candidate pathogenic CNVs were identified (Supplementary Table 4), including an 8.4 Mb deletion of 2q14.1-q14.3 (chr2:116583565-124954598) in one of the two affected individuals in family PK117. Deletions within this region have previously been reported for autism spectrum disorder^65^ and in a patient with a mild holoprosencephaly spectrum phenotype^66^. An overlapping but slightly proximal deletion (~chr2: 114188161-119321989) has been reported to be compatible with a normal phenotype^67^. In one family, PK28, all three affected males in one branch were shown to have a large (6.7 Mb) interstitial duplication spanning cytobands Xp22.31-p22.2.

For homozygous CNVs in HBD regions, we identified homozygous deletions in known ARID genes in several families (Table 3 and Supplementary Table 3), including a 170 kb homozygous deletion spanning 9 of 10 exons of the NS-ARID gene *TUSC3* in family AN21^68^ and a 51 kb deletion spanning exons 37 to 40 of the Cohen syndrome gene, *VPS13B,* in family AN51^69^. Homozygous disrupting loss CNVs included a 50 kb deletion spanning exons 2 and 3 of *RAB8B* in PK95-7; however, this deletion was not in a shared HBD region and was thus considered a potential case of intra-familial genetic heterogeneity.

### Databases for neuropsychiatric disorders

Genes relevant for ID are also frequently identified in individuals with autism spectrum disorders (ASD) and epilepsy (both of which frequently also present with ID). In addition, there is growing support for overlap of ID genes with genes affecting other neuropsychiatric disorders^70^. For example, genes such as *NRXN1* and *ANK3,* which have been linked with SCZ, BD and ASD by CNV and/or genome-wide association studies, are also known ARID genes *(NRXN1:* PTHLS2, MIM 614325; *ANK3:* MRT37, MIM 615494). For this reason, we attempted to evaluate the genes identified here in variant databases or among lists of genes associated with such disorders. We screened various epilepsy or neuropsychiatric disease-specific databases or published datasets for the presence of variants in these genes. In the SCZ/control exome sequence database Genebook^71, 72^, none of the identified variants in Table 2 were present. However, for several of the genes, an increased burden of rare variants has been reported in SCZ cases versus controls (p<0.05 for *VPS35, SYNRG, DMBT1, ALPI,* and *NEU4;* p<0.002 for *SLC13A5),* although when corrected for multiple testing, no individual gene-based tests achieved statistical significance.

In epilepsy databases, we identified *ATRX, MECP2, SLC13A5,* and *ST3GAL3,* all of which have been reported in cases of epilepsy as well as ID. These four genes are either inherited as autosomal recessive or X-linked recessive. *ATRX* and *MECP2* interact with each other and are involved in chromatin binding and gene regulation^73^, and mutations in either gene can have diverse effects on DNA methylation patterns and brain development.

### Protein interaction analysis

We used BioGrid (http://thebiogrid.org) to identify protein interactors for each of the 67 different ARID genes identified among our population (Supplementary Table 3A) as well as *GRIN2B* (Supplementary Table 3C). Interacting proteins are listed, along with gene ontology processes, functions, and cellular compartments, in Supplementary Table 5.

### Pathway analyses

In agreement with past findings of genetic heterogeneity, we observed limited overlap with the gene ontology gene sets. Of the 14,312 sets tested, 16 survived multiple test correction (q-value < 0.05), and a few top-ranked terms are of interest (Supplementary Table 6). Protein glycosylation, which is known to be associated with ID, was ranked 6th with 9 overlapping genes (corrected p < 0.05). As mentioned above, *TPO, DUOX2,* and *TG* are known to be involved in thyroid hormone generation (rank: 14, corrected p < 0.05). The smoothened signalling pathway was ranked 32nd with 4 overlapping genes *(NDST1, CC2D2A, TCTN2,* and *BBS7).*

### Anatomical expression analyses

With the exception of the spinal cord, all neural tissues were enriched in the expression of the ID genes (corrected p < 0.0001, Supplementary Table 8). The frontal cortex was the most enriched, with 1.4 times the expected number of expressed genes. Although the brain expressed a majority of all genes, the ID genes appeared to show specificity. Of the 17 foetal tissues tested, the female gonad and testis had the highest number of expressed genes but did not show enriched expression of the ID genes (corrected p > 0.12).

### Developmental brain expression analyses

Across the developmental transcriptome, ID genes were expressed at higher levels in the normal prenatal brain (Figure 3). The amygdaloid complex showed the most consistent enrichment (5 specimens with significantly higher expression of ID genes). Foetal brain samples from 21 and 24 weeks post-conception showed the highest number of regions with significant expression (> 4 or more). By contrast, enriched expression was not observed in the postnatal brain samples. This prenatal pattern found in exon microarray expression profiles was also observed in RNA sequencing measurements in a largely overlapping set of samples from the same resource (<u>BrainSpan,</u> Supplementary Figure 3,). The RNA sequencing measurements showed donor-specific patterns with many enriched regions. These global patterns were not consistent across donors of the same age, thus suggesting that the RNA sequencing data may have normalization artefacts not present in the microarray measurements. Grouping the ID gene list into genes associated with glycosylation or hormones and metabolism revealed above-average prenatal expression for all groupings (Figure 3). Genes with metabolic or hormonal-associated function had the most stable trajectory. We also observed higher expression for those groups in the prenatal brain in the BrainCloud^74^ resource, which assayed the prefrontal cortex across human development (Supplementary Figure 4).

**Figure 3.**
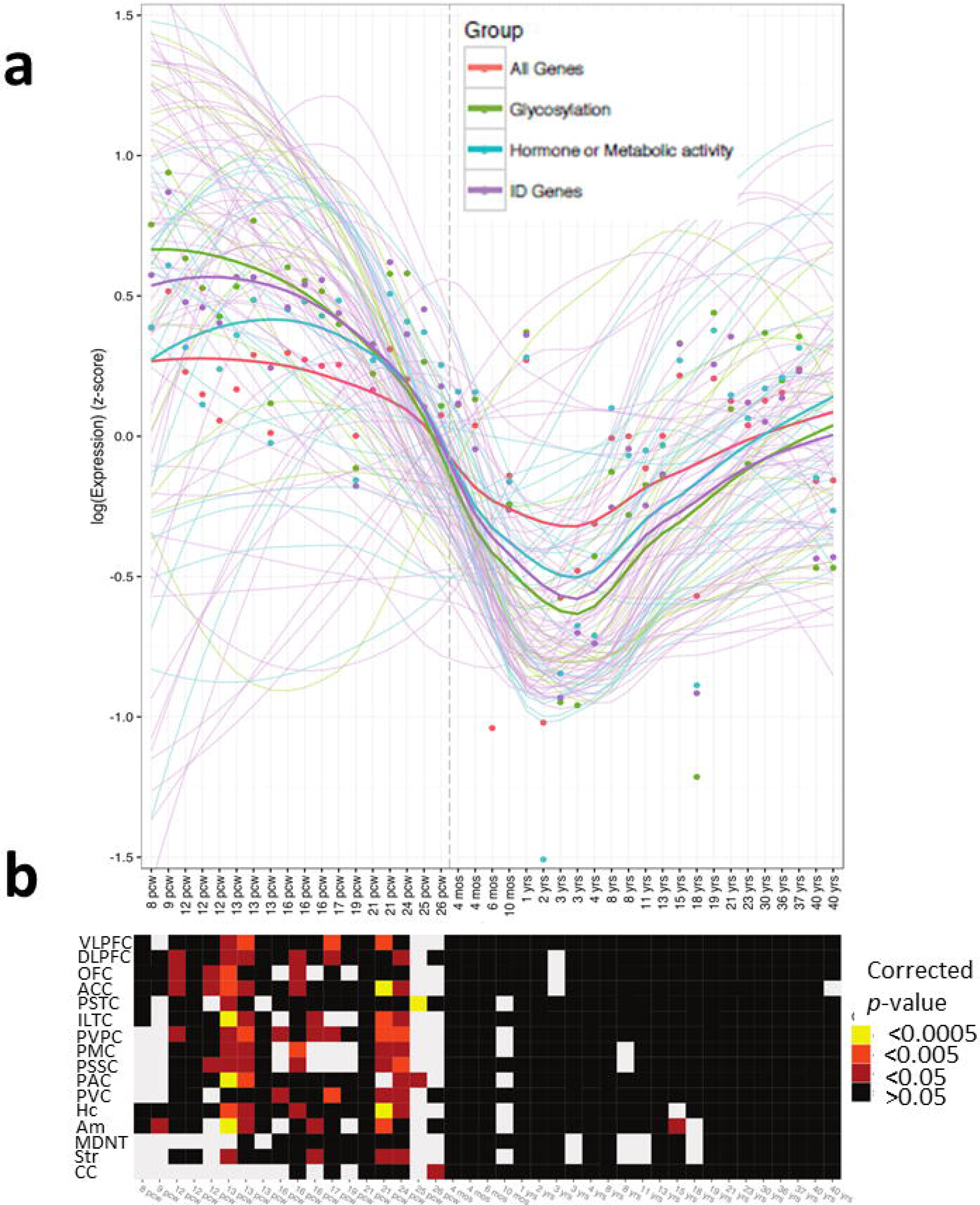
Spatiotemporal expression of ID genes in human brain development: a) Scaled expression trajectories of the ID Genes, averaged across brain regions. Solid lines are average expression for a group of genes. The lighter and thinner lines show trajectories of individual genes. Average expression for each gene group is depicted with points (green: glycosylation-associated (14 genes), blue: hormone or metabolic (20), purple: remainder of the ID Genes (66) and red: remaining genes assayed). Local regression was used to smooth the z-scored expression values for individual genes and gene group averages (LOESS). A vertical dashed line marks birth. **b)** Heatmap showing brain samples enriched for specific expression of ID genes. For each brain region and brain combination, the z-scored expression values for ID genes were compared against all other genes (Wilcoxon one-sided test, FDR-corrected p-values). Significance levels are indicated by black (non-significant result), red (p < 0.05), orange (p < 0.005) and yellow (p < 0.0005). Missing values are shown in grey. Brain regions include the ventrolateral prefrontal cortex (VFC), dorsolateral prefrontal cortex (DFC), orbital frontal cortex (OFC), anterior (rostral) cingulate (medial prefrontal) cortex (MFC), posterior (caudal) superior temporal cortex (area 22c) (STC), inferolateral temporal cortex (area TEv, area 20) (ITC), posteroventral (inferior) parietal cortex (IPC), primary motor cortex (area M1, area 4) (M1C), primary somatosensory cortex (area S1, areas 3,1,2) (S1C), primary auditory cortex (core) (A1C), primary visual cortex (striate cortex, area V1/17) (V1C), hippocampus (hippocampal formation) (HIP), amygdaloid complex (AMY), mediodorsal nucleus of thalamus (MD), striatum (STR) and cerebellar cortex (CBC).

## Discussion

This study describes a cohort of 192 multiplex ID families from Pakistan and Iran. Using combined microarray genotyping, HBD) mapping, CNV analysis, and WES, we identified definite or candidate mutations (or likely pathogenic CNVs) in 51% of families in 72 different genes, including 26 not previously reported for ARID.

Notably, 50% of the variants that we report as single probable mutations were LoF changes. This result compares well with findings from an earlier study reported by Najmabadi et al, 2011^11^, in which 50 new genes for NS-ARID were reported, of which 40% had LoF mutations. In general, LoF mutations provide a higher degree of confidence of disease association than missense mutations.

There are a number of likely reasons that genes/mutations were not found for some families: 1. intra-familial etiologic heterogeneity; 2. poor WES depth of coverage at the etiologic gene/mutation; 3. intronic or intergenic causative mutations, which are not detected by WES. Whole genome sequencing may address this issue; 4. Common variants have been reported in association with many complex diseases, including traits such as intelligence or cognitive ability^75, 76^. It is plausible that a proportion of ID cases with familial aggregation are caused by variants, possibly interacting, that are common in the general population but have low penetrance. These variants together could potentially cause ID; however, this hypothesis has not been explored in ID. 5. Non-genetic factors may be prevalent in some families, e.g., prenatal or perinatal insult^77^.

Given that there may be an overlap in the genetic aetiology of neurodevelopmental and neuropsychiatric disorders, we cross-referenced our ID gene list with those from studies of other neuropsychiatric/neurodevelopmental disorders. Many genes identified for ARID have been implicated across disorders (see Figure 4). For instance, of the newly identified genes, *SLAIN1* has been listed as one of the top-ranked genes for the burden of variants among a large cohort with SCZ (N=1,392)^78^. Data from the Autism Sequencing Consortium^79^ have revealed *de novo* LoF mutations in *SPATA13* and *TBC1D23,* as well as *de novo* missense mutations in *ABI2, TET1,* and *SYNRG.* Li et al (2016)^80^ reported variants in both *MPDZ* and *SPATA13* in ASD cohorts and a *de novo* missense variant in *MAPK8* in SCZ^72^. In addition, whereas heterozygous (typically *de novo)* mutations in *SCN1A* have been linked with ASD, ID, and epileptic encephalopathy, the missense *SCN1A* variant reported here is homozygous. A number of ARID genes (e.g., *NRXN1, CNTNTAP2,* and *ANK3)* have already been implicated in neuropsychiatric disorders by GWAS or CNV studies (as heterozygous). We speculate that, in addition to pleiotropy with other neuropsychiatric/neurodevelopmental disorders, the mode of inheritance may also be variable, (e.g., *SPATA13:* homozygous missense variant in ID (this study) and *de novo* LoF variant in ASD^80^; *HNMT*: homozygous missense variant in ID^17^ and heterozygous splice mutation in SCZ (Genebook)). ID may be conceptualized as a much more severe form of neurodevelopmental disorder than ASD and SCZ.

**Figure 4.**
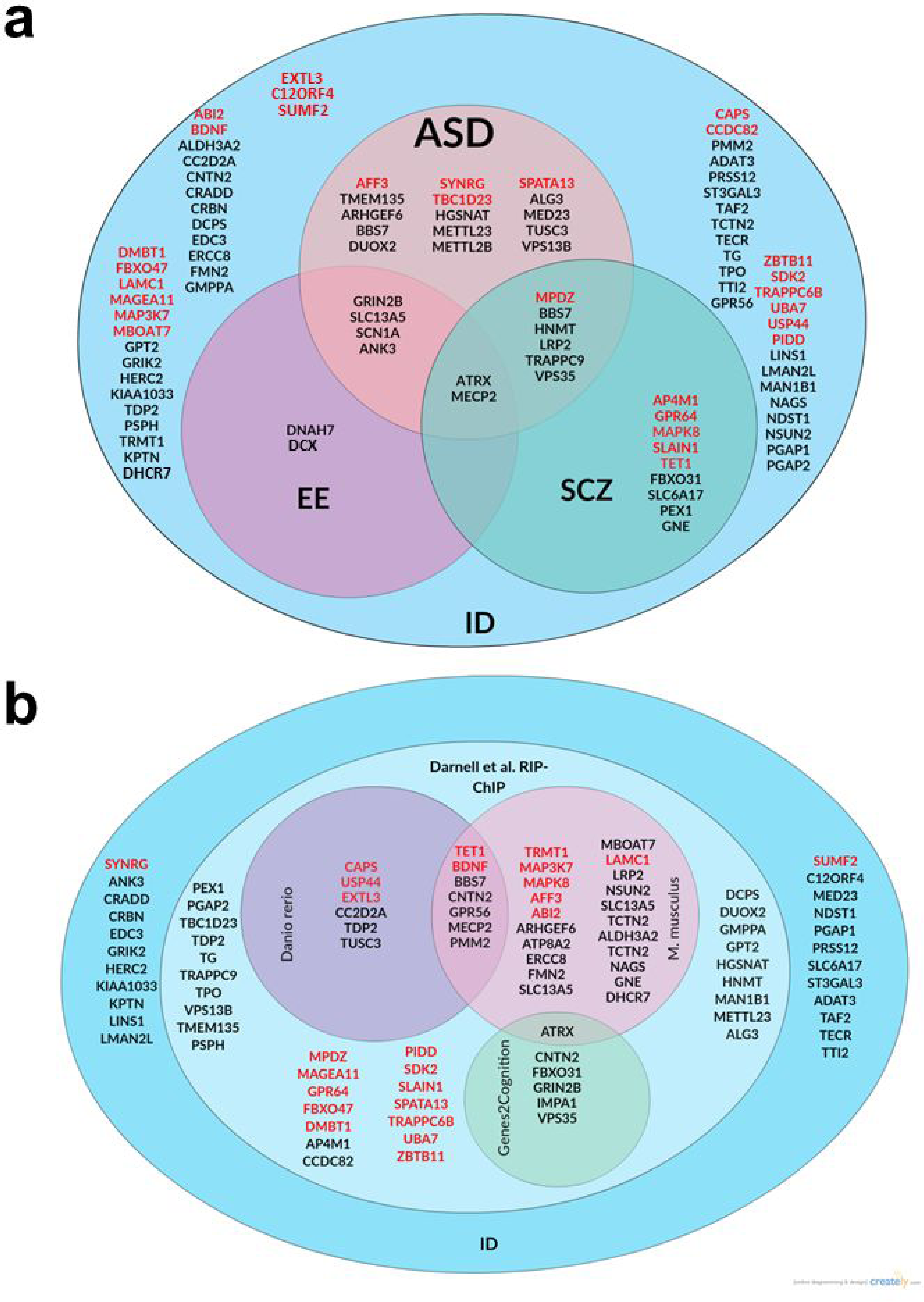
Cross-disorder overlap: The Venn diagrams shown indicate genes for which **a.** variants have been reported in other neuropsychiatric or neurodevelopmental disorders, either homozygous, compound heterozygous, or *de novo,* through searches of published gene lists, including databases such as EpilepsyGene, Gene2Phenotype, Gene2Cognition, Schizophrenia Genebook, and published disease-specific gene sets^6^, HGMD^28^, and OMIM^29^; **b.** functional or animal models with relevant phenotypes have been reported, including resources such as Zfin^30^, targets of FMRP^31^ and Mouseportal (http://www.sanger.ac.uk/science/collaboration/mouse-resource-portal. accessed May 2016). Genes newly reported here are in red text.

Transcriptional analysis of our gene set combined with known ARID genes revealed greater levels of transcription in the prenatal brain than the postnatal or adult brain (Figure 3A) and, in particular, higher levels in the frontal cortex, hippocampus and amygdala (Figure 3B). ID is primarily a disorder of brain development, and thus it was reassuring to observe relevant patterns of spatiotemporal expression of ID genes. Pathway analysis of the gene set showed several significant pathways; thyroid metabolism was prominent, as was protein glycosylation and the Smoothened signalling pathway. This and other studies have predicted involvement of numerous different pathways in ID, which is probably a reflection of the high genetic heterogeneity in ID.

Our findings and those of other groups studying the genetics of NS-ARID should be invaluable for the development of targeted sequencing gene panels for diagnostic screening and for the development of clinical whole exome and whole genome sequencing. In this regard, both the replication of previous findings and new discoveries in this study are important. Replication enhances the confidence in the validity of findings, and new discoveries provide opportunities for further exploration of the functions and biological pathways of the newly associated genes. Cumulatively, such studies have also mapped genes across the human genome in which LoF mutations are viable, and the roles of these genes in human development are thus likely to be amenable to further study and comparison with similar mutations in model organisms. Furthermore, a more comprehensive picture is being assembled of the molecular components and mechanisms that are important for the development of a fully functioning central nervous system, as well as the points in the mechanisms that are most vulnerable to genetic mutation. Through comparison with similar studies, we also note that many more such discoveries are needed to complete the picture. The ultimate challenge, to devise targeted therapeutic strategies for ID patients, is thus a step closer.

The discovery of disease-causing mutations in consanguineous families immediately creates opportunities for carrier screening among relatives and prevention of ID, thus providing a direct benefit to the families, communities, and to public health. In addition, the identification of genetic causes for ID allows ID individuals to be subgrouped on the basis of gene or pathway. This categorization can lead to the development of cohorts that can then be studied prospectively for the natural course of the disease and health complications and that can also be targeted for therapeutic strategies, thus representing a step towards personalized medicine in this important clinical population.

## Acknowledgements

We thank the study participants and their families for their invaluable contributions to this study. We also thank Justin Foong and Andy Wang for their assistance in setting up the NGS analysis pipeline on the CAMH SCC. Computations were performed on the CAMH Specialized Computing Cluster. The SCC is funded by The Canada Foundation for Innovation and the Research Hospital Fund. We also acknowledge the assistance of Muhammad Aslam (LIRD), Tanveer Nasr (Mayo Hospital, Lahore and Chaudhry Hospital, Gujranwala, Pakistan), Muhammad Ilyas (International Islamic University, Islamabad, Pakistan), Reza Najafipour and Soraya Keshavarz (Qazvin University of Medical Science), and Ali Rashidi-Nezhad (Tehran University of Medical Sciences) for family recruitment and phenotyping. We also thank Drs Hans van Bokhoven and Arjen de Brower (Radboud University Medical Center, Nijmegen, The Netherlands), who provided microarray SNP data for families ZA5 and ZA17. RH was supported by a Peterborough K.M. Hunter Charitable Foundation Graduate Scholarship. This study was partially supported by a grant from the Canadian Institutes of Health Research to JBV (#MOP-102758).

## Conflict of Interest

The authors report no conflicts of interest.

Supplementary information is available at the *Molecular Psychiatry* website.

## Supplementary Information

### Supplementary Methods

Supplementary Figure 1: Bioinformatic pipeline for whole exome sequence (WES) analysis.

Supplementary Figure 2: Pedigrees, HomozygosityMapper output and FSuite circo-plots for families with single variants identified, in addition to those shown in Figure 2.

Supplementary Figure 3: Spatiotemporal expression of ID genes in human development using RNA sequencing data.

Supplementary Figure 4: Developmental expression pattern of ID genes in the human prefrontal cortex.

Supplementary Table 1: Family statistics

Supplementary Table 2: Homozygosity-by-descent/autozygosity shared regions, as defined using HomozygosityMapper, cross-referenced with FSuite.

Supplementary Table 3: Mutations identified per family. A. Single homozygous variant identified. B. Two to four variants identified. C. Dominant/de novo mutation identified.

Supplementary Table 4: Pathogenic CNVs and variants of unknown significance identified by microarray analysis.

Supplementary Table 5: BioGRID protein interaction analysis.

Supplementary Table 6: Gene Ontology Pathway analysis

Supplementary Table 7: Gene List for anatomic/temporal transcription analyses.

Supplementary Table 8: Top anatomical regions for ID gene expression.

